## Supplementary Info for "Molecular motor tug-of-war regulates elongasome cell wall synthesis dynamics in *Bacillus subtilis*"

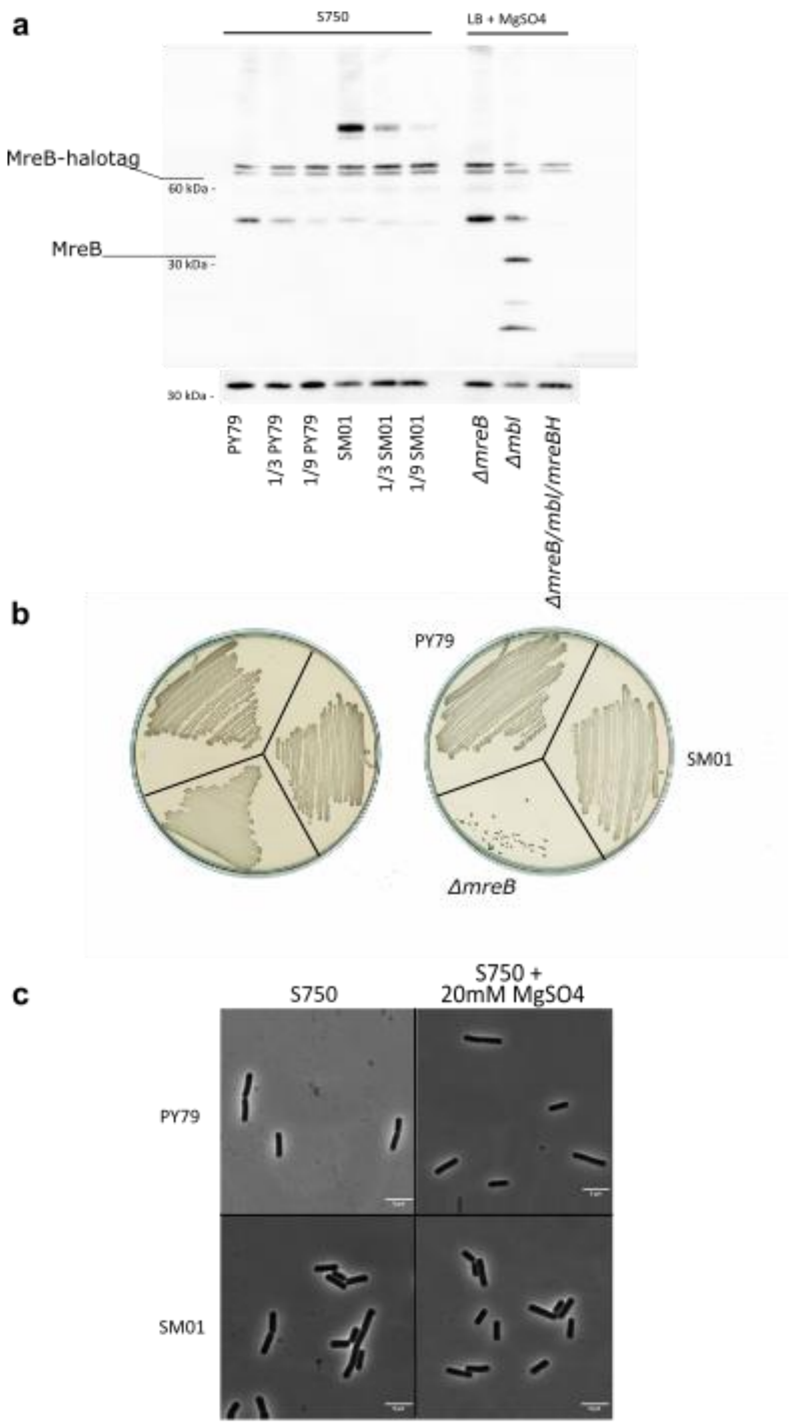

**Supplementary Figure 1. MreB-HaloTag is active but slightly overproduced. (a)** The protein levels of MreB-HaloTag expressed from the native locus (strain SM01) were compared with native levels of MreB in PY79 using western blotting. PY79 and SM01 strains were grown in S750 medium. Please note, other strains were grown in LB supplemented with 20 mM $MgSO_4$ and are used to show the antibody cross activity with Mbl or unspecific binding. MreB was detected using polyclonal MreB antibodies. Detection of Spo0J was used as a loading control. To verify that the signals are not saturated, a serial dilution of the samples was performed. Here the samples were diluted with KS60 (Δ*mreB*, Δ*mbl*, Δ*mreBH*) protein extract. **(b)** MreB-HaloTag can support growth under conditions in which MreB is essential (PAB-Agar plates without added $MgSO_4$), indicating that the fusion protein is active. **(c)** The morphology of PY79 and SM01 is similar in S750 with or without supplemented $MgSO_4$, indicating a well-functioning elongation machinery. Strains used*: B. subtilis* PY79, SM01 (*mreB::mreB-halotag, Δhag*), MB37 (*ΔmreB*), MB35 (*Δmbl*) and KS60 (*ΔmreB, Δmbl, ΔmreBH*).

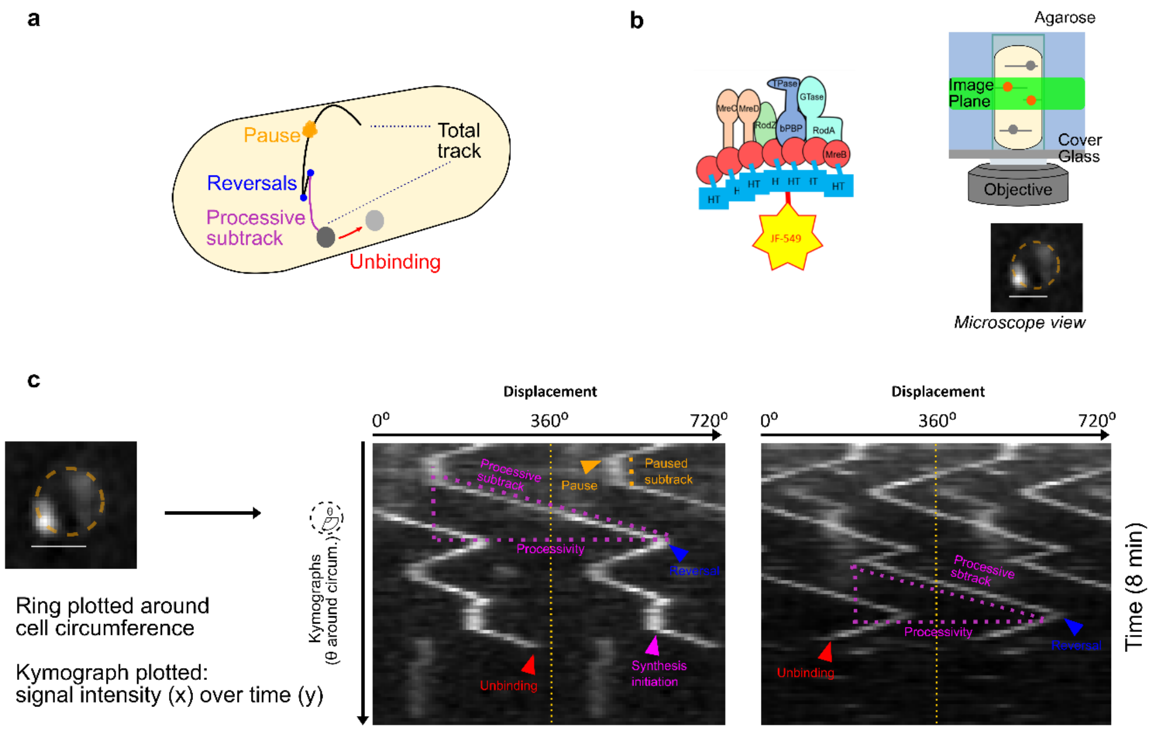

**Supplementary Figure 2. Further description of kymograph acquisition and interpretation. (a)** Schematic illustration showing the MreB dynamics that we observed and measured. **(b)** Cartoon of sm-VerCINI the microscope setup used to acquire single-molecule MreB dynamics and the subsequent view from the microscope where the whole cell circumference can be observed as a circle. **(c)** Example frame of an sm-VerCINI time lapse observing single molecule MreB dynamics. A ring is plotted around the cell circumference and a kymograph is plotted as the fluorescence signal intensity in the x-axis against time in the y-axis. From the kymographs, a variety of MreB dynamics can be identified and quantified as labelled.

To analyse MreB tracking data, kymographs were produced by plotting a ring around the cell circumference based on signal intensity (SI Fig. 2c). The example kymographs (SI Fig. 2c) show MreB displacement around the cell circumference on the x-axis, over time in the y-axis. A diagonal line on the kymograph signifies an MreB molecule moving around the cell circumference and there is a change in both displacement and time, whereas a vertical line shows a static molecule, as there is no displacement around the cell circumference over time.

As we found MreB to be highly processive, with processivity often exceeding one circumference of the cell (360°), it was necessary to plot kymographs over multiple circumferences by repeating the plot multiple times in the x-axis. This is annotated with a dashed yellow line to show each 360° cell circumference. During image analysis, all kymographs were plotted and analysed with 6 repeats of the cell circumference in the x-axis which ensured that no tracks were falsely truncated. The extent at which it was necessary to plot kymographs multiple times around the cell circumference is evident when processivity is plotted as displacement in degrees around the cell circumference (SI Fig. 3).

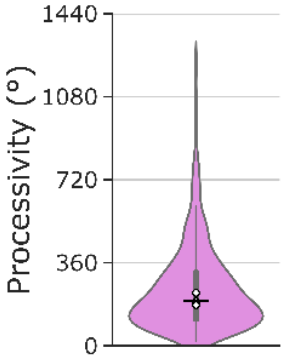

**Supplementary Figure 3.** Processivity of MreB reported in Fig. 1g plotted in degrees around cell short axis.

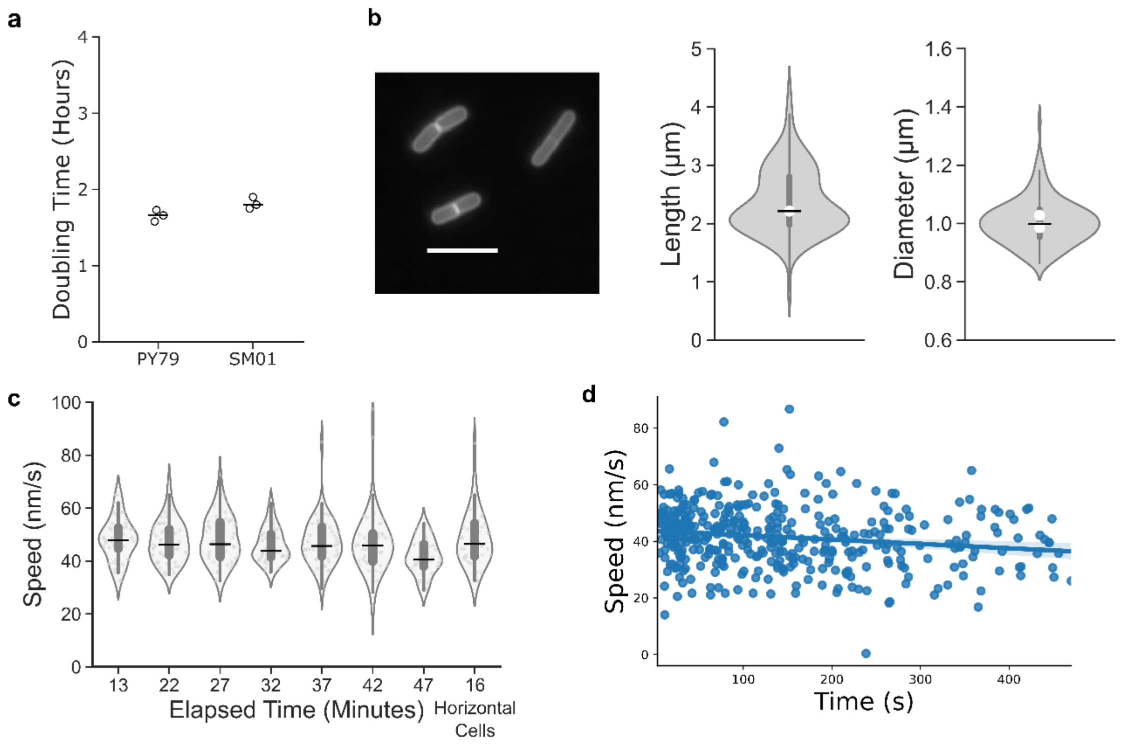

**Supplementary Figure 4: Characterization of *B. subtilis* SM01 (*mreB-HaloTag, Δhag*) growth rate, physiology and MreB-HaloTag speed. (a)** $OD_{600}$ doubling time for SM01 and PY79 (wild type strain). White circles show medians of biological replicates, horizontal lines show overall median while thick vertical lines show IQR. **(b)** Nile Red membrane stain. Scale bar = 5 µm. Violin plots of cell length and diameter. **(c)** Effect of oxygen and/or nutrient depletion on MreB speed over an imaging session. Using single molecule VerCINI, 40 s time-lapse acquisitions with 2 s strobe intervals were taken. **(d)** Representative linear regression control plot used to monitor phototoxicity during an 8-minute time-lapse showing speed of MreB subtracts over the 8-minute acquisition. Blue line shows the linear regression model, while the blue shaded area shows the 95% CI of the model.

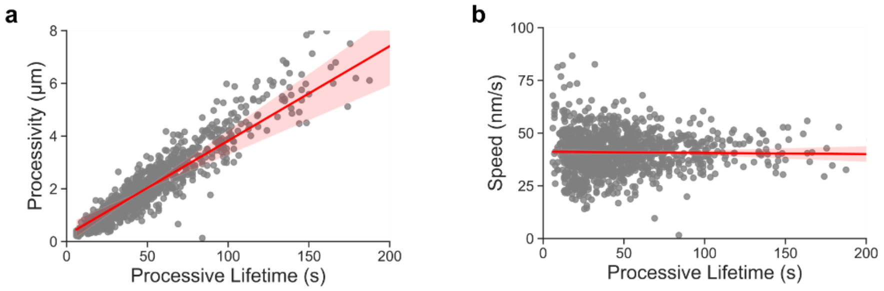

**Supplementary Figure 5: Additional quantification of native MreB dynamics in SM01 (*mreB-halotag, Δhag*) using smVerCINI. (a)** Linear regression plot correlating processive lifetime and processivity ($R^2$ = 0.756252). **(b)** Linear regression plot correlating processive lifetime and speed ($R^2$ = 0.000431). Red lines show the linear regression models, while the red shaded areas show the 95% CI of the models.

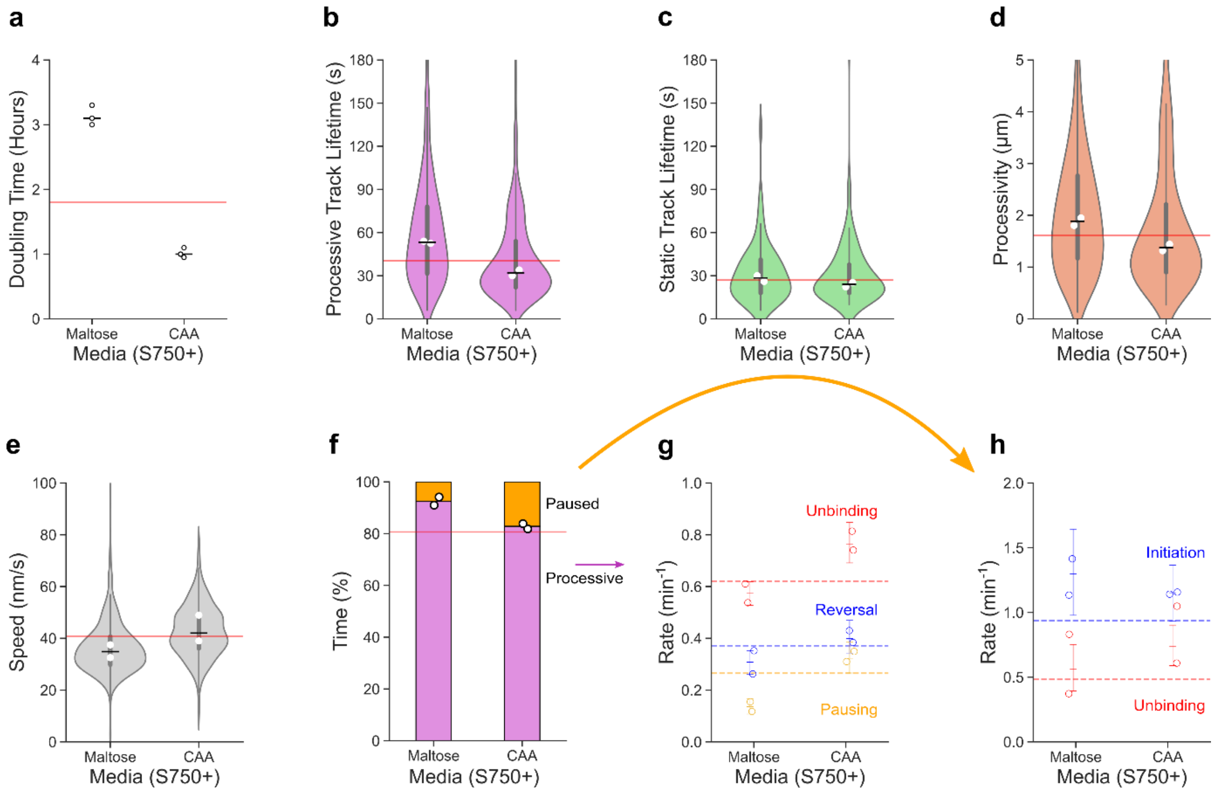

**Supplementary Figure 6: Effect of growth rate on MreB dynamics in strain SM01 (*mreB-halotag, Δhag*). (a)** Doubling time in different growth media. **(b-h)** MreB-HaloTag (JF549) dynamics measured using smVerCINI over 8 minute time-lapse acquisitions in different growth media. **(b-e)** Violin plots of processive and static subtrack lifetime and speed. White circles show medians of biological replicates, horizontal lines show overall median while thick vertical lines show IQR. Red lines show S750$^{glucose}$ medians. **(f-h)** Time in processive and static states and rates of switching from each state. Circles show biological replicate medians. Horizontal lines show S750$^{glucose}$ medians. Error bars show 95% CI.

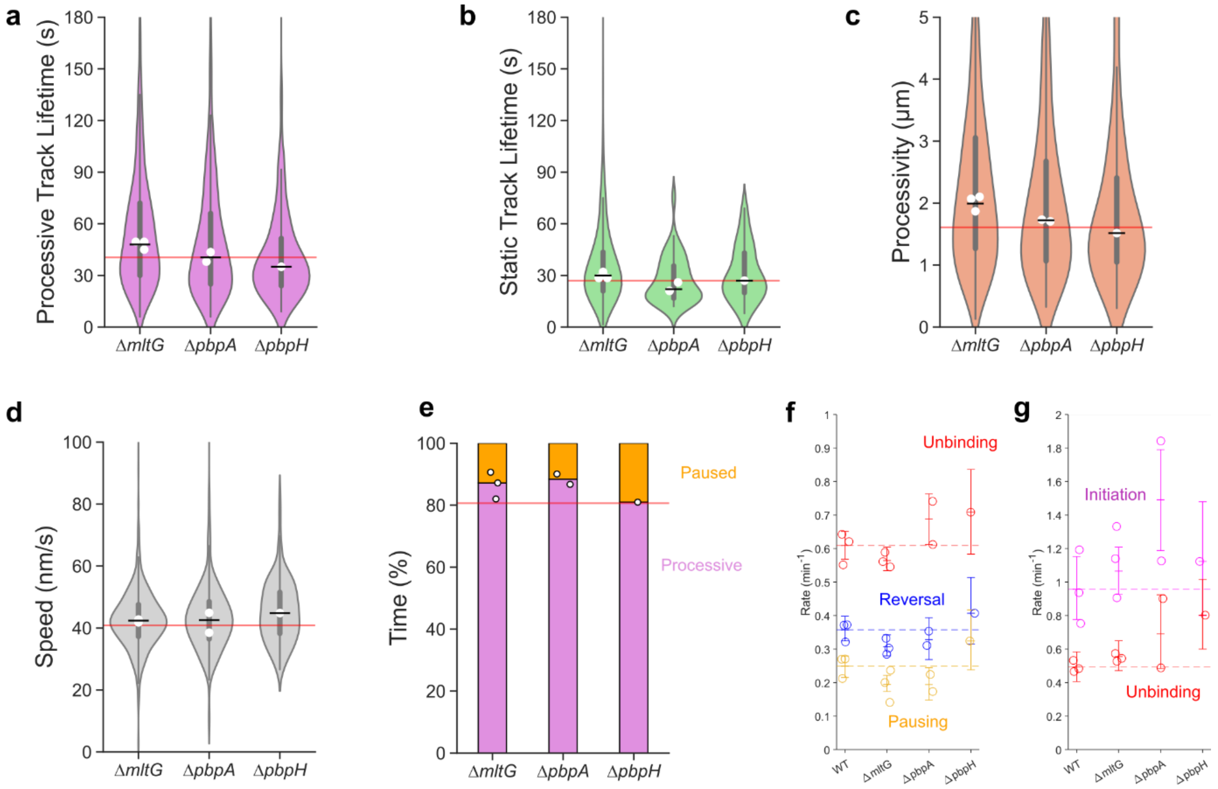

**Supplementary Figure 7: Effect of MltG and aPBP deletion on MreB dynamics. (a-g)** MreB-HaloTag (JF549) dynamics measured using smVerCINI over 8 minute time-lapse acquisitions in strains with the various deletions. **(a-d)** Violin plots processive and static subtrack lifetime, processivity and speed. White circles show medians of biological replicates, horizontal lines show overall median while thick vertical lines show IQR. Red lines show SM01 (*mreB-halotag, Δhag*) medians. **(f-h)** Time in processive and static states and rates of switching from each state. Circles show biological replicate medians. Horizontal lines show SM01 (*mreB-halotag, Δhag*) medians. Error bars show 95% CI. Strains used: SM01 (*mreB-halotag, Δhag*), SM22 (*mreB-halotag, Δhag, ΔpbpA*), SM23 (*mreB-halotag, Δhag, ΔpbpH*), SM41 (*mreB-halotag, Δhag, ΔmltG*).

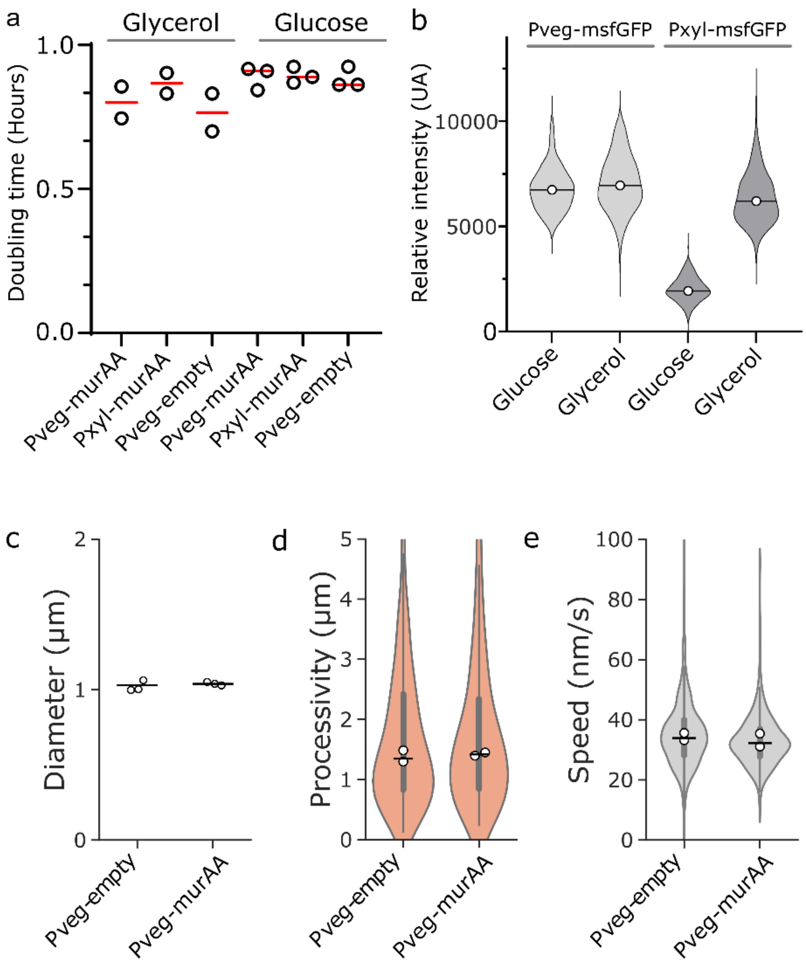

**Supplementary Figure 8. Overexpression of *murAA* does not influence the doubling time, cell width or elongasome processivity or speed in the growth conditions of this study**. $OD_{600}$ **(a)** was measured manually from flask-cultures in S750 glycerol and S750 glucose at 37ºC, showing no difference in growth rate between the strains upon *murAA* overexpression. Expression measurements (b) of the Pxyl and Pveg promoters showed that while Pxyl was not a suitable promoter in our S750 glucose conditions, a constitutive Pveg promoter produced strong protein expression in both S750 glucose and S750 glycerol. This construct was therefore used in our widefield and smVerCINI measurements of murAA overexpression. Cell diameter **(c)** was measured through widefield microscopy, processive velocity **(d)** measured using smVerCINI, and unidirectional processivity **(e)** measured using smVerCINI. Horizontal lines show overall median. Vertical lines indicate IQR. White filled circles represent biological replicates. Microscopy experiments in **c-e** were performed in S750 glucose only as *Bacillus subtilis* cells rapidly stop growing during agarose pad-based microscopy experiments performed in S750 glycerol. Strains used: Bys365 (PY79 *murAA::spec Pxyl-murAA*), MB74 (SM01 *amyE::tet Pveg-murAA*) and MB73 (*SM28 amyE::tet Pveg-empty,* an empty vector control).

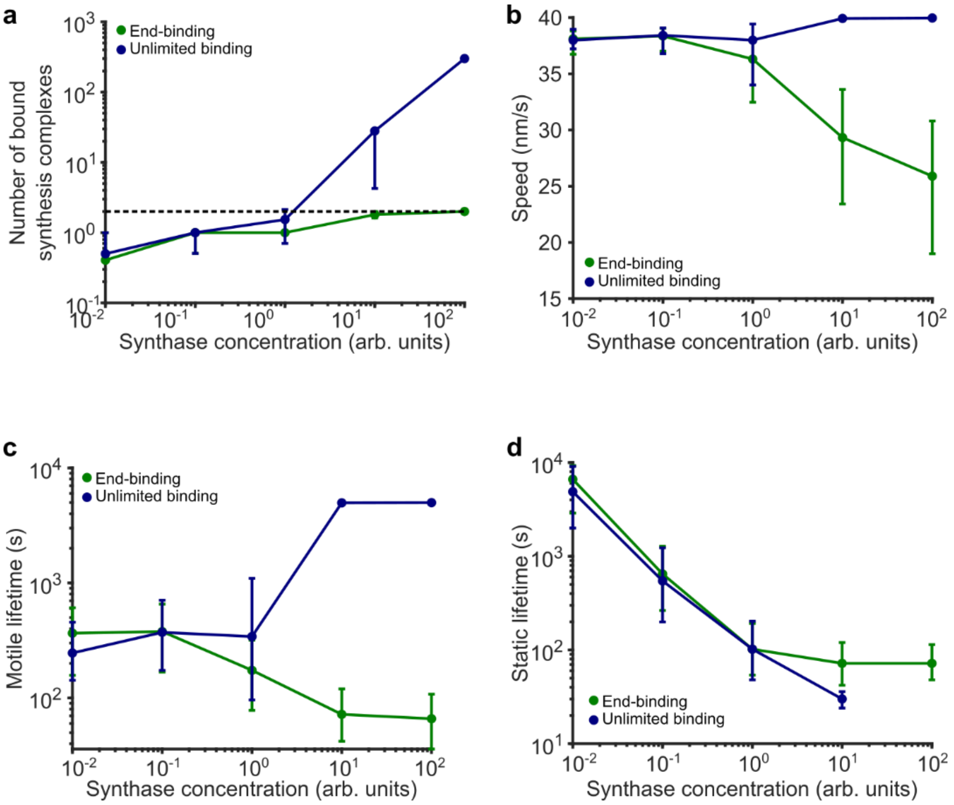

**Supplementary Figure 9: Additional simulation results. (a-d)** Number of bound synthesis complexes, elongasome speed, lifetime of motile elongasome subtracks and lifetime of static elongasome subtracks as a function of synthesis complex concentration for both models. Horizontal dashed line in (a) marks 2 bound synthesis complexes per filament for ease of identification of low synthase concentration regime. Solid coloured lines with filled circles, sample medians; vertical lines, 95% CI.

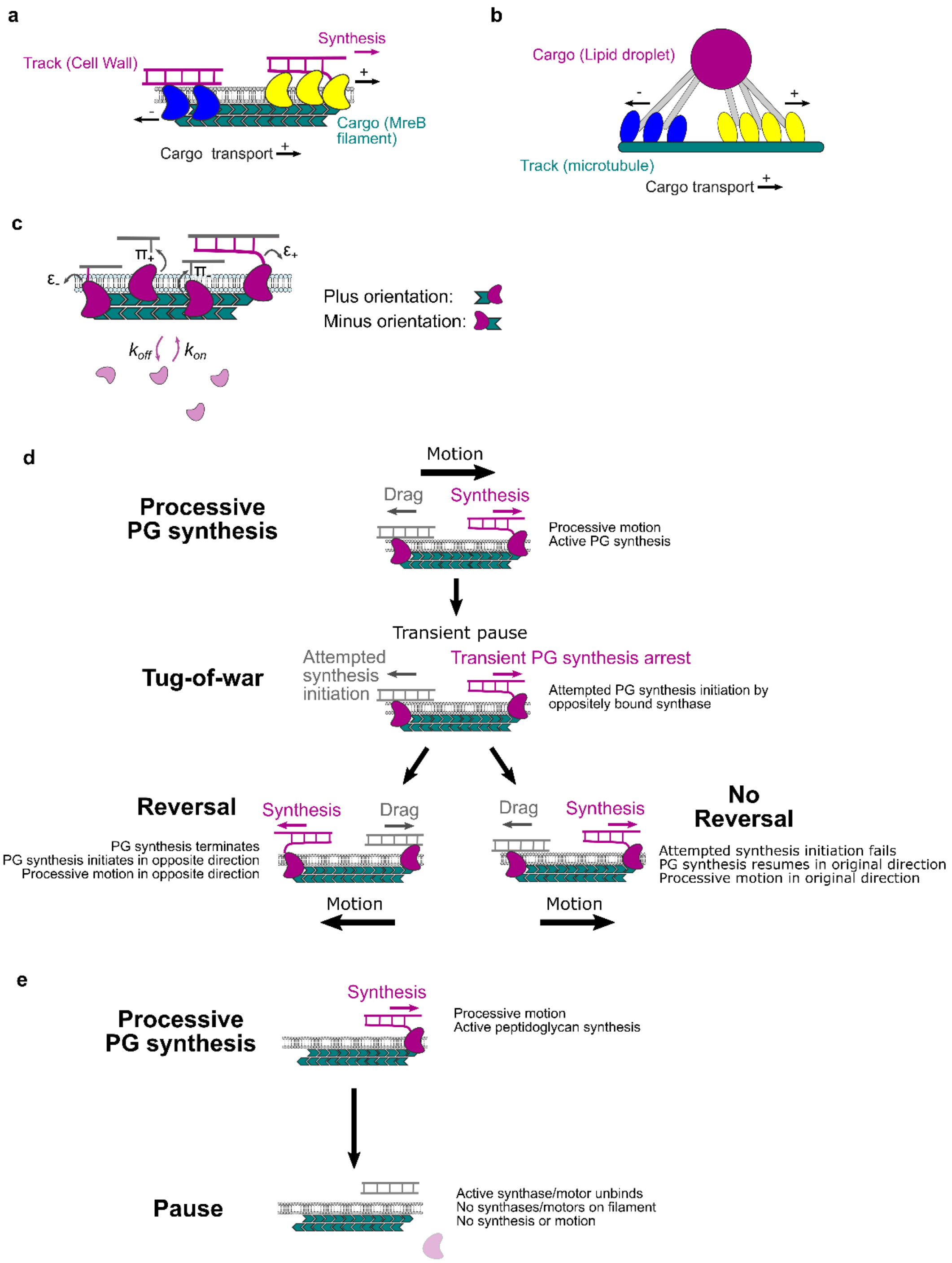

**Supplementary Figure 10. Schematic representations illustrating the differences between molecular tug-of-war models in eukaryotes and the models suggested in the bacterial elongasome. (a)** Proposed elongasome molecular motor tug-of-war model, and **(b)** eukaryotic MKL model for eukaryotic molecular motor tug-of-war from which it is adapted, with colour codes and labels showing how the two models relate to each other [1]. **(c)** Configuration and rates in elongasome tug-of-war model. $k_{on}$/ $k_{off}$, binding/ unbinding rates of synthesis complex (synthase) to MreB filament; π, PG attachment/ initiation rate of MreB-bound synthase; ε, PG detachment/termination rate of MreB-bound synthases actively engaged in PG synthesis. Synthases are assumed able to bind in either a plus or minus orientation to the MreB double filament. **(d)** The mechanisms by which we propose the elongasome reverses by the leading edge becoming the lagging edge of the MreB filament. **(e)** Our proposed mechanism whereby a processively moving elongasome pauses and ceases to actively synthesise peptidoglycan.

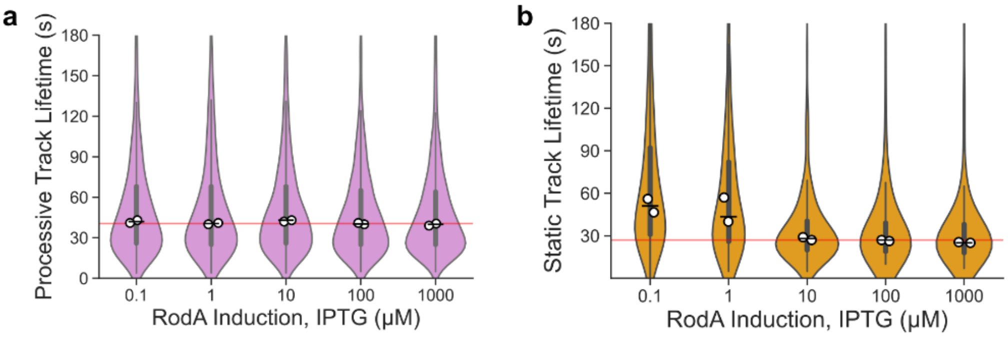

***Supplementary Figure 11.*** **Effect of cellular levels of RodA on track lifetime measured using smVerCINI in strain SM28 (*mreB-HaloTag, Δhag, $P_{spac}$-rodA*) over 8 minute time-lapse acquisitions with various concentrations of IPTG. (a-b)** Violin plots showing processive and static track lifetimes respectively. White circles show medians of biological replicates, horizontal lines show overall median while thick vertical lines show IQR. Red lines show SM01 (*mreB-HaloTag, Δhag*) medians.

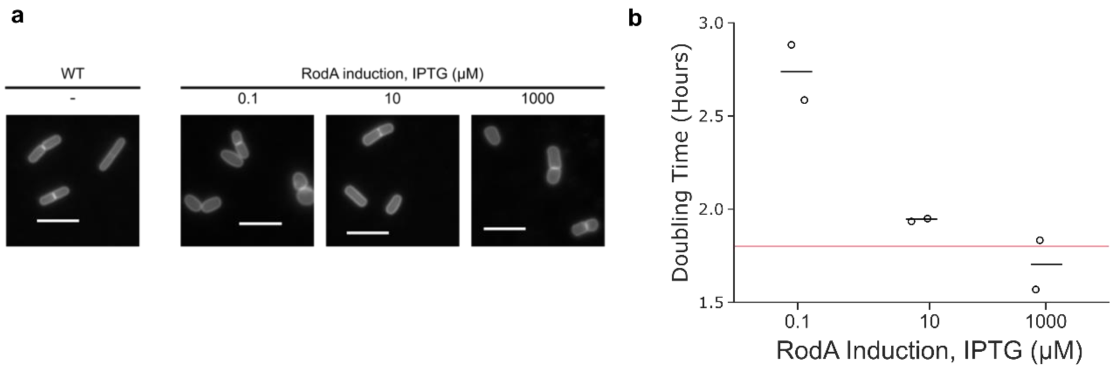

**Supplementary Figure 12. Effect of cellular levels of RodA on cell morphology and doubling time**. **(a)** Membrane stain images of strain SM28 (*mreB-halotag, Δhag,* $P_{spac}$*-rodA*) under various *rodA* induction levels, corresponding to diameter quantification in Fig. 4a. Scale bar = 5 µm. **(b)** $OD_{600}$ doubling time measured manually from flask cultures (as in the microscopy cultures) in strain SM28 (*mreB-halotag Δhag,* $P_{spac}$*-rodA)* under various RodA induction levels and WT strain SM01 (*mreB-halotag, Δhag*) median (red line).

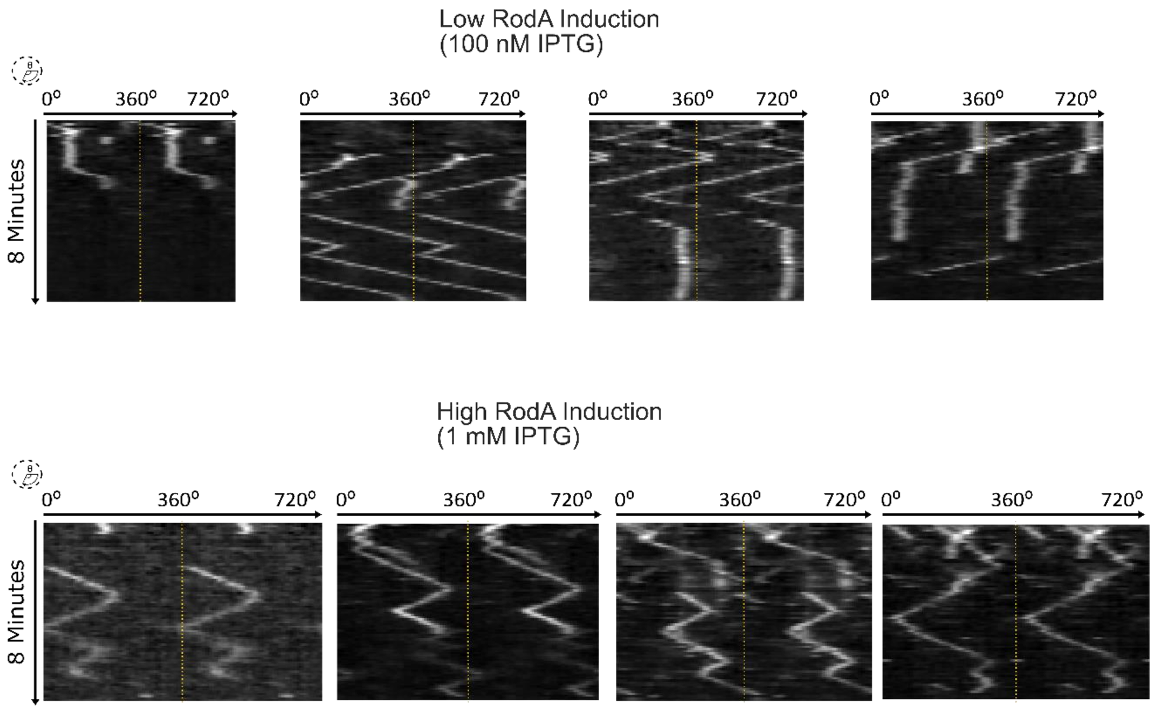

**Supplementary Figure 13.** Further examples of kymographs presented in main text Figure 2.a-b. Kymographs of MreB filament dynamics at low and high RodA levels achieved through expression from an IPTG-inducible promoter. Kymographs are measured around the cell circumference.

**Supplementary Videos 1-3:** Examples of MreB single molecule dynamics in wild-type live B. subtilis cells imaged via smVerCINI corresponding to kymographs Fig 1b. Scale bar = 1 µm.

**Supplementary Videos 4-13:** Examples of MreB single molecule dynamics in low (Supplementary Videos 4-8) and high (Supplementary Videos 9-13) RodA expression level corresponding to kymographs Fig 2a-b and SI Fig 9. Scale bar = 1 µm.

**Supplementary Table 1.** Definitions and descriptions of terms used to describe single molecule trajectories analysed in custom python code in this work.

| Term | Definition/ Description |
|---|---|
| Total Track | A whole observed trajectory. |
| Subtrack | A section of a total track with a constant speed and directionality. |
| Lifetime | The time that a track is observed. Measured as $\Delta y$ on a kymograph. |
| Processive | A subtrack displaying motion around the cell circumference. A subtrack was deemed processive if it travelled $>0.195$ µm. Measured by $\Delta x$ on a kymograph. |
| Paused/Static | A subtrack displaying no motion around the cell circumference. A subtrack was deemed paused/static if it travelled $\leq 0.195$ µm. Measured by $\Delta x$ on a kymograph. |
| Displacement | Distance travelled around the cell circumference. Measured by $\Delta x$ on a kymograph. |
| Processivity | Displacement of a processive subtrack. Measured by $\Delta x$ on a kymograph. The term processivity is used in nucleic acid polymerase fields when describing extension of DNA/RNA, a process physically reminiscent of peptidoglycan synthesis. 'Processivity' was chosen above others such as 'run length', which is often used in association with eukaryotic systems as we think it best describes this particular aspect of elongasome dynamics. |
| Reversal | An event where a processive subtrack changes direction to begin another processive subtrack in the opposing direction. |
| Initiation | An event where a paused subtrack begins motion. |
| Nucleation | An event where the track is observed to appear within the duration of the time-lapse |
| Existing track | A track that is observed from the outset of the time-lapse |
| Pause | An event where a processive subtrack ceases motion to begin a paused/ static subtrack. |
| Unbinding | Loss of signal from a total track. This infers either the molecule unbinding from the cell membrane, or photobleaching of the fluorophore. |
| Switching Rate | Number of each transition type from immobile or processive states, divided by the total duration of all immobile or processive states observed in the dataset [2]. |

**Supplementary Table 2**: **Table of sample sizes.** Shaded areas show quantities not relevant and/or determined; in conditions where *B. subtilis* forms multi-cellular filamentous chains it is not possible to determine the total number of cells used for a measurement without additional cellular labels. Number of biological replicates are experiments performed using independently prepared samples. FOVs refers to the number of fields of view in SIM/ MreB filament density experiments.

| Figure number | Variable | Biological Replicates | Total Tracks | Processive Subtracks | Paused Subtracks | Cells | FOVs |
|---|---|---|---|---|---|---|---|
| 1e | 0.5 s | 2 | 186 | | | | |
| | 2 s | 2 | 274 | | | | |
| | 4 s | 2 | 290 | | | | |
| | 6 s | 3 | 647 | | | | |
| | 8 s | 2 | 361 | | | | |
| 1e-I, SI 2 | Wild-Type dynamics | 3 | 647 | 1078 | 315 | | |
| 2 | 100 nM IPTG | 2 | 1507 | 1696 | 1018 | | |
| | 1 µM IPTG | 2 | 960 | 1286 | 670 | | |
| | 10 µM IPTG | 2 | 1086 | 1669 | 482 | | |
| | 100 µM IPTG | 2 | 821 | 1254 | 302 | | |
| | 1 mM IPTG | 2 | 1431 | 2351 | 495 | | |
| 4a | 100 nM IPTG | 2 | | | | 215 | |
| | 1 µM IPTG | 2 | | | | 166 | |
| | 10 µM IPTG | 2 | | | | 243 | |
| | 100 µM IPTG | 2 | | | | 160 | |
| | 1 mM IPTG | 2 | | | | 261 | |
| 4b | 100 nM IPTG | 2 | | | | | 13 |
| | 10 µM IPTG | 2 | | | | | 17 |
| | 1 mM IPTG | 2 | | | | | 10 |
| SI 4a | - | 2 | | | | | |
| SI 4b | Length | 2 | | | | 119 | |
| | Width | 2 | | | | 153 | |
| SI 6 | Maltose | 2 | 299 | 484 | 75 | | |
| | Glucose + CAA | 2 | 313 | 498 | 140 | | |
| SI 7 | *ΔmltG* | 3 | 747 | 1221 | 295 | | |
| | *ΔpbpA* | 2 | 220 | 343 | 79 | | |
| | *ΔpbpH* | 1 | 119 | 291 | 60 | | |

| | | | | | | |
|---|---|---|---|---|---|---|
| SI 8a | *Pveg-murAA glycerol* | 2 | | | | |
| | *Pxyl-murAA glycerol* | 2 | | | | |
| | *Pveg-empty glycerol* | 2 | | | | |
| | *Pveg-murAA glucose* | 3 | | | | |
| | *Pxyl-murAA glucose* | 3 | | | | |
| | *Pveg-empty glucose* | 3 | | | | |
| SI 8b | *Pveg glucose* | 1 | | | | 340 |
| | *Pveg glycerol* | 1 | | | | 352 |
| | *Pxyl glucose* | 1 | | | | 240 |
| | *Pxyl glycerol* | 1 | | | | 261 |
| SI 8c | *Pveg-empty* | 2 | | | | 538 |
| | *Pveg-murAA* | 2 | | | | 825 |
| SI 8d-e | *Pveg-empty* | 2 | | 948 | | |
| | *Pveg-murAA* | 2 | | 518 | | |
| SI 12 a-b | 100 nM IPTG | 3 | | | | |
| | 1 µM IPTG | 3 | | | | |
| | 10 µM IPTG | 3 | | | | |
| | 100 µM IPTG | 3 | | | | |
| | 1 mM IPTG | 3 | | | | |
| SI 12c | 100 nM IPTG | 3 | | | | |
| | 10 µM IPTG | 3 | | | | |
| | 1 mM IPTG | 3 | | | | |

**Supplementary Table 3a:** Primer list.

| Number | Sequence (5'-3') | Purpose |
|---|---|---|
| oSM01 | ctgcctgcaacaaaagtggtg | Confirm introduction of *hag::erm* into chromosome. |
| oSM02 | gatgtgatctccgcattatcctcac | |
| oSM19 | ctgaattccccctgcgtataatg | Confirm introduction of *pbpA::kan* into chromosome. |
| oSM20 | gaaggggaaaatgaaaccatggaagaag | |
| oSM27 | caaaggtgttacaattaatctcagtatatg | Confirm introduction of *pbpH::kan* into chromosome. |
| oSM28 | gtttaacatgctgcgtatcctgttc | |
| oSM78 | gaatccggtcatcaagctgaaattc | Confirm introduction of *mltG::kan* into chromosome. |
| oSM79 | gtgagctattcccgattgaaactgac | |
| oSM16 | gtcatatttcgtgtagctgaaaaag | Confirm introduction of *rodA':kan-P$_{spac}$-rodA* into chromosome. |
| oSM43 | gttgcgtaaaagaagaagaatacccac | |
| MB-F-62 | ataaatacaggtgttatattattaaacgagaaaggagattcctagg atgagcaaaggagaagaacttt | Amplify msfGFP to build pMB10. |
| MB-F-63 | tcacattaattgcgttgcgcttatttgtagagctcatccatg | |
| MB-F-64 | cgtttaataatataacacctgtatttattta | Amplify pCW433 to build pMB10. |
| MB-F-65 | gcgcaacgcaattaatgtga | |
| MB-F-66 | agaaaggagattcctaggatgatggaaaaaatcatcgtccgc | Amplify *murAA* to build pM12. |
| MB-F-67 | tcacattaattgcgttgcgcttatgcatttaagtcagaaacga | |
| MB-F-68 | catcctaggaatctcctttctcg | Amplify pCW433 to build pMB13. |
| MB-F-69 | aataaatacaggtgttatattattaaacg *gcgcaacgcaattaatgtga* | |

**Supplementary Table 3b:** Strain list

| Strain | Species and Strain | Relevant Genotype | Construction |
|---|---|---|---|
| bYS40 | *B. subtilis PY79* | *mreB::mreB-halotag* | Published strain [3]. Provided by Ethan Garner Lab. |
| - | *B. subtilis 168* | *trpC2 hag::erm* | BKE library [4]. |
| SM01 | *B. subtilis PY79* | *mreB::mreB-halotag hag::erm* | bYS40 transformed with *hag::*erm gDNA. |
| - | *B. subtilis 168* | *trpC2 pbpA::kan* | BKK library [4]. |
| SM22 | *B. subtilis PY79* | *mreB::mreB-halotag hag::erm pbpA::kan* | SM01 transformed with *pbpA::kan* gDNA. |
| - | *B. subtilis 168* | *trpC2 pbpH::kan* | BKK library [4]. |
| SM23 | *B. subtilis PY79* | *mreB::mreB-halotag hag::erm pbpH::kan* | SM01 transformed with *pbpH::kan* gDNA. |
| YK2245 | *B. subtilis 168CA* | *trpC2 rodA':kan-*$P_{spac}$*-rodA* | Published strain [5]. Provided by Richard Daniel Lab. |
| SM28 | *B. subtilis PY79* | *mreB::mreB-halotag hag::erm rodA':kan-*$P_{spac}$*-rodA* | SM01 transformed with YK2245 gDNA. |
| - | *B. subtilis 168* | *trpC2 mltG::kan* | BKK library [4]. |
| SM41 | *B. subtilis PY79* | *mreB::mreB-halotag hag::erm mltG::kan* | SM01 transformed with *mltG::kan* gDNA. |
| MB60 | *B. subtilis PY79* | *SM28, amyE::tet pveg-murAA* | SM28 transformed with pMB12. |
| MB59 | *B. subtilis PY79* | *SM28, amyE::tet pveg-empty* | SM28 transformed with pMB13. |
| MB36 | *B. subtilis PY79* | *SM28, murAA::spec pxyl-murAA* | SM28 transformed with murAA:: spec pXyl-murAA gDNA. |
| MB38 | *B. subtilis PY79* | *PY79, amyE::tet Pveg-msfGFP* | PY79 transformed with pMB10 |
| MB76 | *B. subtilis PY79* | *PY79, amyE::spec Pxyl-msfGFP* | PY79 transformed with amyE::spec Pxyl-msfGFP gDNA. |
| MB37 | *B. subtilis PY79* | *PY79, ΔmreB::neo* | PY79 transformed with mreB::neo gDNA. |

| | | | |
|---|---|---|---|
| MB35 | *B. subtilis PY79* | *PY79, Δmbl::zeo* | PY79 transformed with *mbl::zeo* gDNA. |

**Supplementary Table 4.** Equipment used to test PureDenoise.

| | Equipment 1: (2020 PC) | Equipment 2: (2019 Laptop) | Equipment 3: (2017 PC) |
|---|---|---|---|
| **System** | Linux | Mac OS | Windows 10 |
| **CPU** | Intel Core i7 10700K (16) | Intel Core i9 9880H (8) | Intel Core i7-7700K |
| **GPU** | NVIDIA 3090 | AMD Readon 5500M | NVIDIA 1080TI |

**Supplementary Table 5.** Speed test for various PureDenoise settings.

| Settings | 1024*1024pixels*300frames, 4 cycle spin, 3 adjacent frames (seconds) | Average (seconds) |
|---|---|---|
| GPU: NVIDIA 3090 | 137.18/116.81/117.45/118.50/121.90 | 122.36 |
| GPU: NVIDIA 1080TI | 228.30/229.42/246.00/229.75/230.13 | 232.72 |
| GPU: AMD Readon 5500M | 292.51/283.15/276.64/274.37/284.88 | 282.31 |
| CPU: Intel Core i7 10700K (with 1 thread) | 2689.34 | 2689.34 |
| CPU: Intel Core i9 9880H (with 1 thread) | 3618.33 | 3618.33 |
| CPU: Intel Core i7-7700K (with 1 thread) | 2967.69 | 2967.69 |
| CPU: Intel Core i7 10700K (with 4 thread) | 965.24 | 965.24 |
| CPU: Intel Core i9 9880H (with 4 thread) | 1042.22 | 1042.22 |
| CPU: Intel Core i7-7700K (with 4 thread) | 1096.30 | 1096.30 |
| CPU: Intel Core i7 10700K (with 16 thread) | 755.87/743.24/748.26 | 749.12 |
| CPU: Intel Core i9 9880H (with 16 thread) | 698.33/735.26/700.48 | 711.35 |
| CPU: Intel Core i7-7700K (with 16 thread) | 1069.37/1056.00/1061.04 | 1062.13 |

***Supplementary Table 6:*** Effect sizes between experimental conditions

| **Figure number** | **Quantity, comparison conditions** | **Difference** | **95% CI** | ***No of data points (N1, N2)*** | ***Data type*** | **Number of biological replicates** |
|---|---|---|---|---|---|---|
| 2d | *Unbinding rate,* RodA induction: "1 mM" minus "100 nM" | 0.12 $min^{-1}$ | [-0.03,0.06] | 1696,2351 | Processive sub-tracks | 2 |
| | *Reversal rate,* RodA induction: "1 mM" minus "100 nM" | 0.20 $min^{-1}$ | [0.17,0.24] | 1696,2351 | Processive sub-tracks | 2 |
| | *Pausing rate,* RodA induction: "1 mM" minus "100 nM" | -0.13 $min^{-1}$ | [-0.17,-0.10] | 1696,2351 | Processive sub-tracks | 2 |
| 2e | *Initiation rate,* RodA induction: "1 mM" minus "100 nM" | 0.66 $min^{-1}$ | [0.50,0.81] | 1018,495 | Static sub-tracks | 2 |
| | *Unbinding rate,* RodA induction: "1 mM" minus "100 nM" | 0.14 $min^{-1}$ | [0.06,0.23] | 1018,495 | Static sub-tracks | 2 |
| 2f | *Speed,* RodA induction, "1 mM" minus "100 nM" | -19.76 nm/s | [-20,-19] | 1696,2351 | Processive sub-tracks | 2 |
| 2g | *Processivity,* RodA induction, "1 mM" minus "100 nM" | -0.91 µm | [-1.0,-0.78] | 1696,2351 | Processive sub-tracks | 2 |
| 4a | *Cell diameter, RodA induction, "1 mM" minus "10 µM"* | 1.08 µm | [0.92, 1.3] | 243, 215 | Cells | 2 |
| | *Cell diameter, RodA induction, "100 nM" minus "10 µM"* | 1.50 µm | [1.3, 1.7] | 243, 262 | Cells | 2 |
| 4b | *Processive MreB filament density, RodA induction, "1 mM" minus "10 µM"* | 0.36 track density ($µm^{-2}$) | [-0.54,1.3] | 17, 10 | Fields of view | 2 |
| | *Processive MreB filament density, RodA induction, "100 nM" minus "10 µM"* | -3.2 track density ($µm^{-2}$) | [-4.2,-2.3] | 17, 13 | Fields of view | 2 |
| SI 6d | *Processivity,* "Maltose" | -0.51 µm | [-0.68,-0.35] | 484,498 | Processive sub-tracks | 2 |

| | | | | | | |
|---|---|---|---|---|---|---|
| | minus "Glucose+CAA" | | | | | |
| SI 7c | Processivity, *ΔmltG* minus WT | 0.24 µm | [0.16, 0.33] | 1222 | Processive sub-tracks | 3 |
| SI 7d | Speed, *ΔmltG* minus WT | 0.18 nm/s | [0.10, 0.27] | 1222 | Processive sub-tracks | 3 |
| SI 7c | Processivity, *ΔpbpA* minus WT | 0.01 µm | [-0.05, 0.19] | 343 | Processive sub-tracks | 2 |
| SI 7d | Speed, *ΔpbpA* minus WT | 0.24 nm/s | [0.12, 0.37] | 343 | Processive sub-tracks | 2 |
| SI 7c | Processivity, *ΔpbpH* minus WT | -0.06 µm | [-0.20, 0.11] | 191 | Processive sub-tracks | 1 |
| SI 7d | Speed, *ΔpbpH* minus WT | 0.44 nm/s | [0.29, 0.60] | 191 | Processive sub-tracks | 1 |
| Si 8a | Doubling time, "*Pveg-murAA*" minus "*Pveg-empty*" | 0.035 | [-0.13,0.20] | 2,2 | Growth rates | 2 |
| | Doubling time, "*Pxyl-murAA*" minus "*Pveg-empty*" | 0.10 | [-0.045,0.25] | 2,2 | Growth rates | 2 |
| | Doubling time, "*Pveg-murAA*" minus "*Pveg-empty*" | 0.007 | [-0.068,0.082] | 2,2 | Growth rates | 2 |
| | Doubling time, "*Pxyl-murAA*" minus "*Pveg-empty*" | 0.011 | [-0.052,0.074] | 2,2 | Growth rates | 2 |
| SI 8c | Diameter, "*Pveg-murAA*" minus "*Pveg-empty*" | 15 | [-73,104] | 825,538 | Cells | 2 |
| SI 8d | Processivity, "*Pveg-murAA*" minus "*Pveg-empty*" | 0.022 | [-0.081,0.13] | 518,948 | Processive sub-tracks | 2 |
| SI 8e | Speed, "*Pveg-murAA*" minus "*Pveg-empty*" | -0.19 | [-0.29,-0.090] | 518,948 | Processive sub-tracks | 2 |

**Supplementary note 1.** Technical details on the simulation.

We implemented stochastic simulations of elongasome tug-of-war, based on the Müller, Klumpp, and Lipowsky (MKL) model of tug-of-war in eukaryotic cargo transport[1]. The key differences between the elongasome tug-of-war model presented here and MKL model are that:

1. Whereas MKL assume fixed number of motors, we extend the model to allow stochastic concentration dependent motor (synthesis complex) binding and unbinding from the cargo (MreB filament).
2. We limit the analysis to identical motors (synthesis complexes) binding in each plus/ minus orientation to a cargo, here the MreB filament, instead of allowing the possibility of different types of motors.
3. We model two cases: *end-binding*, where a maximum of two motors can be bound, one in the plus orientation and one in the minus orientation (ie at opposite ends of the MreB filament); and *unlimited binding,* where unlimited motors can bind in each orientation.

We adapt the terminology of the model for peptidoglycan synthesis rather than cargo transport, a model schematic with key rates included as labels is shown in Supplementary Figure 6. Comparing the MKL and elongasome models, elongasome peptidoglycan synthesis complexes (hereafter synthases, for brevity) correspond to molecular motors, MreB filaments correspond to the cargo. Most importantly, where MKL refer to motor binding/ unbinding to microtubules and initiation/ termination of processive transport, we consider instead synthase attachment/ detachment from the cell wall, leading to initiation or termination of PG synthesis.

We describe here the key features of the elongasome tug-of-war model. The equations for MreB filament (cargo) force and speed are unchanged compared to the MKL model, so those results are presented without proof. Derivations of force and speed equations can be found in the supplement of the study describing the MKL model [1] and further discussion of the model in a review by Bresslof and Newby [6].

We simulate a membrane bound MreB filament, from which active synthesis complexes can bind with concentration dependent rate  and constant unbinding rate:

$$k_{on} = k_{on0}[Synthase],$$

$$k_{off} = Const.$$

Synthases can bind to the filament in two possible orientations, plus or minus, with equal rates for each binding orientation. The total number of synthases bound in each direction, $N_+, N_-$, respectively is randomly determined by the binding/ unbinding rates.

In the single isolated synthase case, MreB-bound synthases stochastically attach to the cell wall and initiate synthesis at constant attachment rate $\pi_0$, and terminate synthesis and detach from the cell wall at detachment rate $\varepsilon_0$. Synthases are modelled to move with load dependent velocity $v(F)$, driven by peptidoglycan synthesis:

$$v(F) = \begin{cases} v_F(1 - F/F_s) \text{ for } F \leq F_s \\ v_B(1 - F/F_s) \text{ for } F \geq F_s \end{cases}$$

Here $v_F$ is the speed of an isolated processive synthase (set as 40 nm/ s based on experiments). $F$ is the force arising from tug-of-war with oppositely bound synthases. $F_s$ is the stall force required to arrest forward motion. $v_B$ is the backwards stepping synthase speed, which is assumed negligible for a polymerization driven motor, but is set to just above zero (0.1 nm/ s) to avoid singularities in the model.

The PG synthesis termination/ detachment rate is assumed to increase exponentially with applied force $F$, based on Kramers rate theory*:*

$$\varepsilon(F) = \varepsilon_0 \exp(|F|/F_d)$$

When multiple synthases are allowed to bind to the MreB filament and interact with the cell wall, a tug-of-war scenario is encountered. The number of active synthases performing/ initiating PG synthesis at a given time in the plus direction is $n_+$. The number of active synthases in the minus direction is $n_-$. The number of active synthases in each direction is determined by the constant synthase attachment/ synthesis initiation rate $\pi_0$, and the force dependent detachment/ synthesis termination rates $\varepsilon(F)$.

In the case of more active plus synthases, $n_+ > n_-$, the MreB filament moves in the plus direction and the filament (cargo) force and velocity are

$$F_c(n_+, n_-) = \lambda n_+ F_s + (1 - \lambda)n_- F_s,$$

$$v_c(n_+, n_-) = \frac{n_+ - n_-}{n_+/v_F + n_-/v_B},$$

$$\text{where } \lambda = 1/(1 + {}^{n_+ v_B}/_{n_- v_F}).$$

In the case $n_- > n_+$, $v_B$ and $v_F$ are swapped in the above equations and the MreB filament moves in the minus direction. If no synthases are engaged in PG synthesis, the force and MreB filament speed are both zero.

In the elongasome tug-of-war model, as in the original MKL model, when one plus synthase is engaged in PG synthesis, and a minus motor initiates PG synthesis, the forces on each motor are equal, so the resulting tug of war is equally likely to result in either MreB filament reversal, or failed minus-end initiation and continued plus end synthesis. However, when multiple synthases are simultaneously engaged in plus end synthesis, successful minus-end synthase initiation and MreB filament reversal becomes exponentially less likely. This is because the force on the single initiating minus end synthase is $n_+$ times larger than the force on each plus end synthase, $F_- = n_+ F_+$, and because PG synthesis termination rate/ detachment rate depends exponentially on the force on the motor:

$$\varepsilon_+ = \varepsilon_0 \exp\left({}^{F_c}/_{n_+ F_d}\right)$$

$$\varepsilon_- = \varepsilon_0 \exp\left({}^{F_c}/_{n_- F_d}\right)$$

Stochastic simulations of MreB filament dynamics using the above model were performed using the Gillespie algorithm.

*Choice of simulation parameters.*

As noted previously, the strong motor scenario in the MKL model, where the detachment force $F_d$ is far lower than the stall force $F_s$, reliably generates tug-of-war-driven reversals [1]. This case would correspond to the biological situation where minus-end synthases attempt attach to the PG, and initiate synthesis, forcibly terminating synthesis of synthases engaged in plus-end synthesis. This scenario seems most likely to correspond to a plausible model of elongasome tug of war and was thus chosen for further investigation, with Fd and Fs arbitrarily chosen as 1 pN and 100 pN respectively, as it is the ratio of Fd/Fs which affects simulation dynamics rather than the absolute value.

The rate of synthase binding to MreB scales with concentration, at low simulated concentration the basal synthase binding rate $kOn_0$ was set to the observed rate of initiation from static to motile MreB filaments at low RodA concentration, 0.5 $min^{-1}$ (Figure 2E). This could correspond to either the binding of an entire synthesis complex (eg RodA-PBP2A-MreCD) to the MreB filament, or binding of one rate limiting protein, such as RodA, to an otherwise assembled synthesis complex already bound to the MreB filament.

The PG synthesis initiation rate $\pi_0$ was assumed to be more frequent and set to $10kOn_0$ higher, which promotes frequent tug of war based on the phase space described for the original MKL model [1]. The basal rate of PG synthesis termination $\varepsilon_0$ was set to the observed MreB pausing rate at low RodA concentration 0.3 $min^{-1}$ (Figure 2D). Synthase unbinding from the complex - or more precisely the unbinding of one or more components such as RodA, PBP2A or PBP2H required for formation of a complete active synthesis complex - was assumed to occur rapidly after synthesis termination, based on the observation that each of these three proteins have substantial diffusive populations, in addition to the elongasome associated processive populations [7] so the unbinding rate $kOff_0$ was set to $10\varepsilon_0$.

Two possible synthase models were tested, *end-binding* and *unlimited-binding*. In the *end-binding* model, synthases can only bind to the ends of the asymmetric MreB double filament, in a single orientation, so the maximum number of plus or minus end bound synthases is one, and maximally two motors (one plus and one minus) can be bound at a given time, with the average number of bound motors depending on the simulated concentration. In the *unlimited-binding* model, synthases can bind all along the filament in either orientation, so there are no limits on the number of plus or minus bound motors, which is determined only by the simulated concentration.

The simulated concentration affects the synthase binding rate $kOn_0[Synthase]$. We simulated a range of concentrations approximately corresponding to experimental observed MreB motility rates.

Full source code for stochastic model and simulation scripts are available on the Holden lab GitHub: https://github.com/HoldenLab/lipowskiModel

**Supplementary Note 2. S750 media preparation protocol.**

S750 was prepared fresh for every experiment as we found that using older (over ~2 weeks) media led to substantial experimental variability, and in particular led to reduced MreB speeds. See supplementary table 5 for S750 media and stock component recipes. 100 X metal mix was prepared from powder and stock solutions no earlier than 6 months before use. Individual metal stock solutions were prepared no earlier than 1 year before use and stored at 4 °C. S750 salts were from powder prepared no earlier than 2 months before use and stored at 4 °C. Metal mix and S750 salts were stored at 4 °C. S750 media was prepared in a 100 ml volume within 2 days of every experiment and stored at room temperature in a dark cupboard. S750 salts and metal mix were sterilised through a 0.2 µm filter. Glutamate, MiliQ water and carbon sources were sterilised by autoclave and stored at room temperature.

**Supplementary Table 7**. Composition of S750 media and supplements used in this work. All % units are weight to volume

| Medium/ Solution | Composition/ Concentration/ Vendor |
|---|---|
| Metal Mix (100 X) | 2 mM Hydrochloric Acid (HCl) (Honeywell)<br><br>190 mM Magnesium Chloride Hexahydrate (Sigma)- From powder<br><br>65.9 mM Calcium Chloride Dihydrate (Sigma- From powder<br><br>4.84 mM Manganese Chloride Tetrahydrate (Fisher)- From 100 mg/ml stock solution<br><br>0.106 mM Zinc Chloride (Sigma)- From 10 mg/ml stock solution<br><br>0.196 mM Thiamine Chloride (Sigma)- From 10 mg/ml stock solution in 2 mM HCl<br><br>0.470 mM Iron (III) Chloride Hexahydrate (VWR)- From 10 mg/ml stock |
| S750 Salts (10 X) | 500 mM MOPS (Sigma)<br><br>100 mM Ammonium Sulphate (Sigma)<br><br>Potassium Phosphate Monobasic (VWR)<br><br>pH adjusted to 7.0 with Potassium Hydroxide (VWR) |
| S750 Media | 1 X S750 salts (Above)<br><br>1 X Metal mix (Above)<br><br>1 % Carbon source (Glucose or Maltose (VWR))<br><br>10 mM L-Glutamate (Sigma)<br><br>Made to 100 ml volume with MiliQ water |

### REFERENCES

[1] M.J.I. Müller, S. Klumpp, R. Lipowsky, Tug-of-war as a cooperative mechanism for bidirectional cargo transport by molecular motors, Proc. Natl. Acad. Sci. 105 (2008) 4609–4614. https://doi.org/10.1073/pnas.0706825105.

[2] G. Özbaykal, E. Wollrab, F. Simon, A. Vigouroux, B. Cordier, A. Aristov, T. Chaze, M. Matondo, S. van Teeffelen, The transpeptidase PBP2 governs initial localization and activity of the major cell-wall synthesis machinery in E. coli, eLife 9 (2020) e50629. https://doi.org/10.7554/eLife.50629.

[3] S. Hussain, C.N. Wivagg, P. Szwedziak, F. Wong, K. Schaefer, T. Izoré, L.D. Renner, M.J. Holmes, Y. Sun, A.W. Bisson-Filho, S. Walker, A. Amir, J. Löwe, E.C. Garner, MreB filaments align along greatest principal membrane curvature to orient cell wall synthesis, eLife 7 (2018) e32471. https://doi.org/10.7554/eLife.32471.

[4] B.-M. Koo, G. Kritikos, J.D. Farelli, H. Todor, K. Tong, H. Kimsey, I. Wapinski, M. Galardini, A. Cabal, J.M. Peters, A.-B. Hachmann, D.Z. Rudner, K.N. Allen, A. Typas, C.A. Gross, Construction and Analysis of Two Genome-scale Deletion Libraries for Bacillus subtilis, Cell Syst. 4 (2017) 291-305.e7. https://doi.org/10.1016/j.cels.2016.12.013.

[5] K. Emami, A. Guyet, Y. Kawai, J. Devi, L.J. Wu, N. Allenby, R.A. Daniel, J. Errington, RodA as the missing glycosyltransferase in Bacillus subtilis and antibiotic discovery for the peptidoglycan polymerase pathway, Nat. Microbiol. 2 (2017) 16253. https://doi.org/10.1038/nmicrobiol.2016.253.

[6] P.C. Bressloff, J.M. Newby, Stochastic models of intracellular transport, Rev. Mod. Phys. 85 (2013) 135–196. https://doi.org/10.1103/RevModPhys.85.135.

[7] J. Domínguez-Escobar, A. Chastanet, A.H. Crevenna, V. Fromion, R. Wedlich-Söldner, R. Carballido-López, Processive Movement of MreB-Associated Cell Wall Biosynthetic Complexes in Bacteria, Science 333 (2011) 225–228. https://doi.org/10.1126/science.1203466.
